# A leaf-carbon economy model to explain the adaptive value of low vs. high leaf mass per area as a function of habitat productivity

**DOI:** 10.64898/2026.11.09.765326

**Authors:** Ilan Elhanan, Efrat Sheffer

**Affiliations:** Institute of Plant Sciences and Genetics in Agriculture, The Robert H Smith Faculty of Agriculture, Food and Environment, The Hebrew University of Jerusalem. 229 Hertzel street, Rehovot, Israel 7610001

**Keywords:** functional traits, LMA, leaf lifespan, plant ecology, Specific leaf area, maximum growth rate, non-equilibrium model, interest rate, carbon investment

## Abstract

Plant traits interact with environmental conditions to determine plant fitness. In particular, plant species with low leaf mass per area (LMA) are more common in high fertility soils and forest gaps, while plant species with high LMA are more common in low fertility soils and in shaded forest understories. It is not yet known why the adaptive value of LMA changes as a function of environmental productivity. We present a model for plant growth as a function of LMA and productivity. Our model relies on an empirically derived relationship between LMA and leaf lifespan. We are able to explain why species with lower LMA are successful in high productivity environments including environments with high availability of either light or soil nutrients, by showing that the LMA which maximizes growth rate decreases with increasing rate of carbon assimilation. We show that low LMA species are better at taking advantage of the ‘high interest’ environment of high productivity habitats by reinvesting assimilated carbon in new leaves and earning compound interest, while high LMA species grow more efficiently in the ‘low interest’ environment of low productivity habitats. This tradeoff sheds light on the adaptive value of LMA as it relates to environmental productivity.

## Introduction

Plant functional traits are used to find general rules governing the interactions between plant species and their environment (McGill et al., 2006, Westoby and Wright, 2006, Adler et al., 2013, Moles, 2018). A key trait is leaf mass per area (LMA), which is correlated with many other leaf traits such as leaf nitrogen concentration, leaf phosphorus concentration, maximum rate of net photosynthesis per leaf mass (but not per leaf area: Osnas et al., 2013) and leaf lifespan (Wright et al., 2004, Wright et al., 2005). This set of correlated leaf traits gives rise to what is termed the *leaf economics spectrum* (Wright et al., 2004).

In particular, LMA is consistently correlated with leaf lifespan (Poorter et al., 2009). For example, the LMA and leaf lifespan of shade-tolerant trees is high compared to the low LMA and short leaf lifespan of shade-intolerant trees (Baltzer and Thomas, 2010, Kitajima and Poorter, 2010, Laurans et al., 2012). Similarly, the LMA and leaf lifespan of plant species that inhabit nutrient-poor soils is high compared to the low LMA and leaf lifespan of plants that inhabit nutrient-rich soils (Reich et al., 2004, and showing the pattern for LMA only: Read et al., 2005, Werner and Homeier, 2015, Jager et al., 2015).

A well supported explanation for the correlation between LMA and leaf lifespan is that leaves with high LMA contain more mechanical support tissues (of different types; Castro-Díez et al., 2000, Shipley et al., 2006, Poorter et al., 2009, Kitajima et al., 2012, Onoda et al., 2017) compared to low LMA leaves. Mechanical support tissues increase the resilience of the leaves to a range of environmental hazards (like herbivory and snow) and therefore result in a longer leaf lifespan compared to leaves with less mechanical support (Poorter et al., 2009).

The mechanical support explanation provides a mechanism for the consistent correlation between LMA and leaf lifespan, but not an explanation for their ecological distributions. Why do plant species from low-productivity niches (e.g. shaded forests) or environments (e.g. nutrient-poor soil) have more resilient leaves compared to plant species from high productivity niches (e.g. canopy gaps) or environments (e.g. nutrient-rich soils)? Conservation of nutrients (nitrogen, phosphorus) in the plant was suggested as the main explanation for the high resilience (and long lifespan) of leaves of plant species from low productivity environments compared to the low resilience (and short lifespan) of leaves of plant species from high productivity environments (Aerts and Chapin III, 1999, Weng et al., 2017).

The nutrient conservation hypothesis provides an explanation for the patterns of LMA and leaf lifespan across soil-productivity gradients (Weng et al., 2017). However, nutrient conservation cannot directly explain the patterns of LMA and leaf lifespan across regeneration niches (e.g., shade-tolerant vs. shade-intolerant species).

There are also models that predict that species with low LMA values will perform better in high light environment while species with high LMA values will perform better in the shade (Sterck et al., 2006, Falster et al., 2018). These models explain patterns of LMA and leaf lifespan with respect to light environment but cannot explain the patterns of LMA and leaf lifespan across soil-productivity gradients.

We propose a hypothesis that can explain patterns of LMA and leaf lifespan across both soil productivity gradients and regeneration niches. Our hypothesis is based on carbon economy (e.g. Falster et al. 2018) without needing to explicitly consider other nutrients. We postulate that low LMA species are better at taking advantage of ‘high interest’ environments by reinvesting assimilated carbon in new leaves and earning compound interest on that carbon investment, while high LMA species are more efficient in ‘low interest’ environments due to their higher leaf lifespan and therefore longer period of carbon revenue stream. The ‘interest rate’ of the environment is determined by the rate of assimilation, which is higher in high productivity habitats than in low productivity habitats. Our model can therefore explain patterns of LMA and leaf lifespan with respect to any productivity gradient.

## Methods

We modeled the carbon economy of plants to analyze how plant growth is influenced by LMA and leaf lifespan. We chose to model plants during their early period of growth. During early growth, leaf area can expand quickly without causing excessive self-shading, which allows plants to receive a fast return on investment when investing the carbon assimilated by leaves to expand further their leaf area. The early period of plant growth is critical for plant establishment and therefore influences plant fitness and persistence throughout its life cycle. Our model applies only for plants during this early period of intensive growth, it does not apply for mature plants at equilibrium of leaf growth and leaf senescence.

We analyzed the model to look for the LMA value and the corresponding leaf lifespan value that optimizes plant growth in different environments and explored the implications of our model in the context of the distribution of plant species with different values of LMA across productivity gradients.

**Table 1:** Definitions and units of the variables and parameters of the model.

| Symbol | Unit | Description |
|---|---|---|
| ***B*** | *g* | Plant biomass |
| ***LL*** | *days* | Leaf lifespan |
| ***LMA*** | *g* · *m*<sup>−2</sup> | Leaf mass per area |
| ***NAR*** | *g* · *days*<sup>−1</sup> · *m*<sup>−2</sup> | Net assimilation rate per leaf area |
| ***LMF*** |  | Leaf mass fraction |
| ***α*** |  | Slope of the log-log relationship of LMA and leaf life span |
| ***β*** |  | Intercept of the log-log relationship of LMA and leaf life span |
| ***γ*** | *days*<sup>−1</sup> | Turnover rate for non-leaf tissue |
| $\boldsymbol{\frac{\partial B}{\partial t}}$ | *g* · *days*<sup>−1</sup> | Biomass growth rate |
| ***c*** |  | Efficiency of turning assimilates to biomass |

We assume that LMA and leaf lifespan (***LL***) are mechanically related, such that plants with higher LMA values have longer leaf lifespan in the form of equation 1.

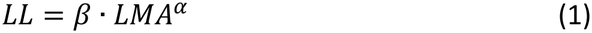

This form is based on empirical evidence (Wright et al., 2004), where according to empirical evidence *α* > 1, *β* > 0, and both are constants (Wright et al., 2004, Falster et al., 2012). We also explore the meaning of this empirically-based assumption using values of 0 < *α* ≤ 1, to examine how this can change the predictions of the model.

The net growth rate of plant biomass (***B***) is the outcome of the rate at which biomass is acquired (***B***_***gain***_) minus the rate at which biomass is lost (***B***_***loss***_):

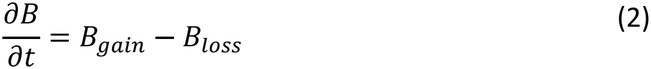

Biomass is lost when old plant tissues (root, stem and leaf) senesce and are shed. The rate at which leaf biomass (***LB***) is being lost is an inverse of leaf lifespan (eq. 3), and so, the higher the LMA the lower the rate of leaf loss. Using the relationship between leaf lifespan and LMA, we get that the rate at which leaf biomass is lost is:

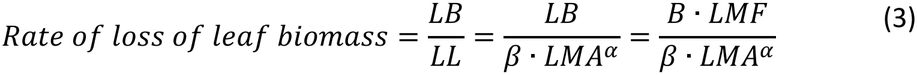

where ***LMF*** is leaf mass fraction, the ratio of leaf biomass to total plant biomass, which we assume for simplicity to be independent of other leaf traits (this assumption is non-essential, see supplementary material for details). We add a term for the loss of non-leaf tissue (root and stem) to include all biomass loss in the form:

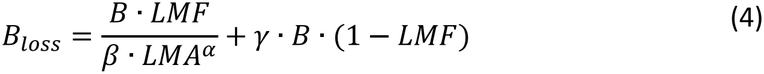

where *γ* represents the average rate of biomass loss of non-leaf tissue. We model biomass gain through carbon capture in photosynthesis and the consequent use of carbon in the construction of new plant biomass, as follows:

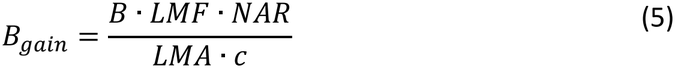

The term *B* · *LMF* represents leaf biomass, divided by *LMA* it gives the total leaf area of the plant. *NSR* represents the net assimilation rate (assimilation minus respiration) of carbon per unit leaf area. We assume that *NSR* is independent of LMA. While the rate of photosynthesis per unit mass was found to be positively correlated with LMA, the rate of photosynthesis per unit of leaf area was found to be independent of LMA (Osnas et al., 2013). The constant ***c*** represents the average construction cost for new plant biomass, that is, the efficiency at which assimilated carbon is transformed to plant biomass. Putting everything together we get at:

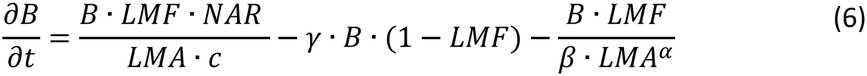

Eq. 6 shows a tradeoff between the rate at which carbon is acquired and the rate at which carbon is lost, as both biomass loss and biomass gain are inversely related to LMA. Plants with higher LMA values will have a lower rate of carbon gain but also a lower rate of carbon loss.

## Results

A numerical solution of our model (eq. 6) shows that for each value of net assimilation rate (***NAR***), there is one value of LMA that optimizes the relative growth rate (Fig. 1). The value of LMA that optimizes growth rate decreases with increasing NAR. Changes in all other parameters influence the values of relative growth rate and the values of optimal LMA, but still maintain a single optimal LMA value.

**Figure 1:**
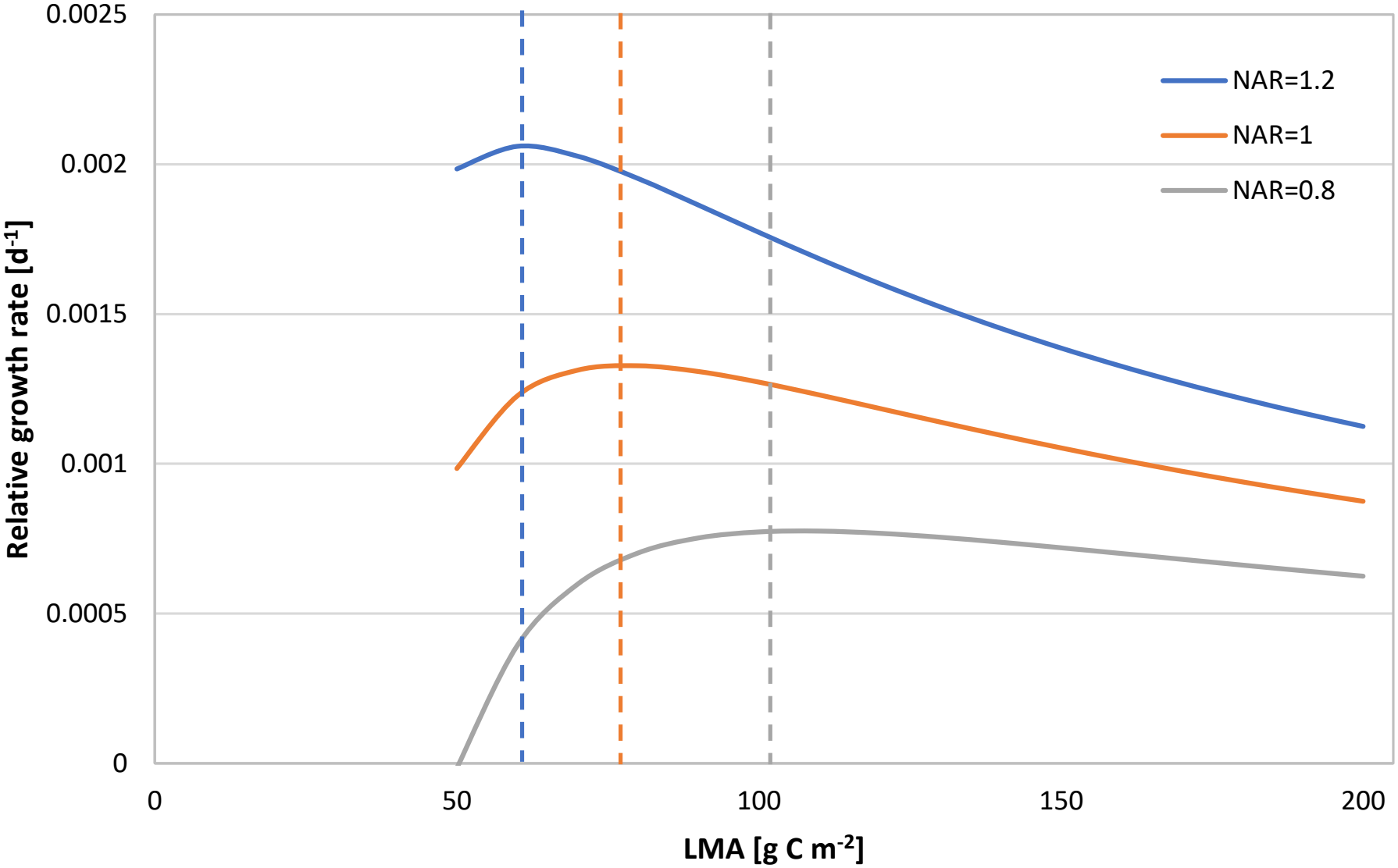
Daily rate of relative growth as a function of leaf mass per area (LMA) with three different values of the net assimilation rate (NAR). Dashed lines mark the LMA at which growth rate is maximal for each curve. The values of all other parameters are as follows: *α* = 1.71, *β* = 0.092923, c = 1.2, LMF = 0.3 and *γ* = 0. The values of *α* and *β* are based on Wright et al., 2004, the values of c and LMF are typical values for these traits and *γ* was arbitrarily set to zero since it does not influence the shape of the curve.

An analytical solution of the model (see Supplementary material) also shows that a single LMA value optimizes the growth rate. The LMA value that optimizes the growth rate is given by:

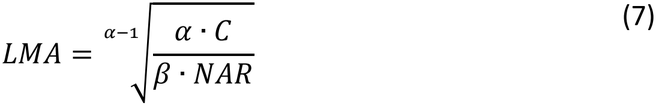

The LMA value that optimizes relative growth rate decreases as a function of increasing assimilation rate ***NAR*** (eq. 7), as we also demonstrate numerically in Fig. 2. Assuming that NAR increases with productivity (as discussed below), we show that the optimal LMA is expected to decrease with increased productivity of the environment. Therefore, our model analysis predicts that species with low LMA values will have a higher relative growth rate than species with high LMA values when growing in high productivity environments. Vice versa, species with high LMA values will have a higher relative growth rate than species with low LMA values when growing in low productivity environments

**Figure 2:**
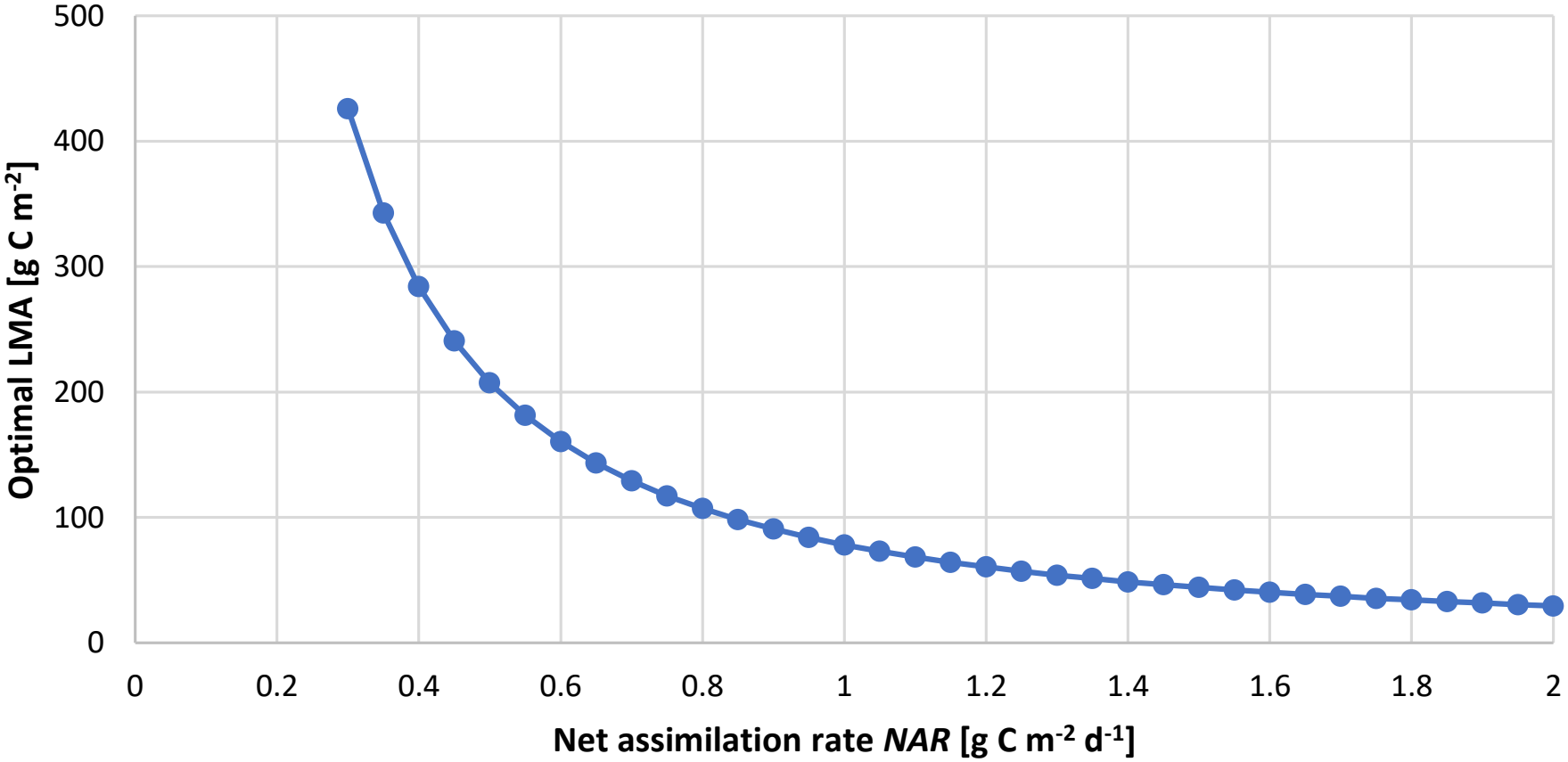
Value of LMA that optimizes growth rate as a function of net assimilation rate NAR. This figure was created using the same parameters as figure 1.

The conclusion that there is a single finite value of LMA that optimizes growth rate requires the condition of *α* >1. Otherwise, when 0 < *α* ≤ 1 no such finite value of LMA exists, and growth rate either keeps increasing as LMA tends to infinity, or keeps increasing as LMA tends to zero, or is constant (see further elaboration in the Supplementary material).

## Discussion

We presented a model in which the LMA value that optimizes growth rate depends on the net assimilation rate per unit of leaf area. To infer the implications of this finding for the relationship between LMA and productivity, we first explore the relationship between NAR and productivity. We assume that in all plants NAR increases with increasing soil fertility. In particular, we assume NAR is positively correlated with nitrogen availability because enzymes involved in photosynthesis require large amounts of nitrogen, and increased nitrogen availability leads to increased number (and size) of chloroplasts per leaf area and therefore also to a higher rate of assimilation per unit area (Lawlor and Tezara, 2009). We also assume that NAR increases with increasing light availability (at least in shaded environments), as higher irradiation directly leads to higher rates of assimilation (up to a limit). Finally, high availability of water should also lead to high NAR because it allows stomata to remain open thus increasing assimilation rates. To summarize, an increase in productivity via an increase in the availability of plant resources (light, water, or soil nutrients) is expected to increase NAR.

The LMA value that optimizes the relative growth rate decreases with increasing net assimilation rate. The relative growth rate is one of the components of fitness (albeit not the only one). Therefore, based on our model we predict that plants with low LMA values will have a higher fitness in high productivity habitats (with high NAR) compared to plants with high LMA values. Therefore, plants with low LMA values will be selected in high productivity habitats, while plants with high LMA values will be selected in low productivity habitats (with low NAR). These findings coincide with the available evidence. Shade-intolerant species have lower LMA than shade-tolerant species (Baltzer and Thomas, 2010, Kitajima and Poorter, 2010, Laurans et al., 2012), and species from fertile soils have lower LMA than species from infertile soils (Reich et al., 2004, Read et al., 2005, Werner and Homeier, 2015, Jager et al., 2015). We acknowledge that NAR may also be affected by other (endogenous) plant traits. Therefore, the LMA value that optimizes growth can vary among species as a function of other traits that influence NAR in addition to the (exogenous) effect of environmental conditions.

Our conclusion is also supported by other, more specific models. A model for carbon gain in closed canopy conditions (i.e., equilibrium), revealed that the LMA value that maximizes carbon gain decreases with increasing soil nitrogen availability (Weng et al., 2017). This model is similar to our model in the sense that both assume a positive relationship between LMA and leaf lifespan, but Weng et al. (2017) specifically model the nitrogen economy of the plant therefore focusing on the nutrient conservation hypothesis while our model provides a general framework for the relationship between LMA and productivity.

Sterck et al. (2006) also support a similar conclusion that low LMA species grow faster than high LMA species in canopy gap conditions (high productivity), while high LMA species are better than low LMA species in the shaded understory (low productivity). Their model of the carbon economy and growth rate of shade-tolerant vs. shade-intolerant species uses real data of leaf traits from a specific set of species and therefore implicitly includes the relationship between LMA and leaf lifespan. Therefore, Sterck et al. (2006) can be seen as a private case of our model.

Falster et al. (2018) analyzed a model that arrived at a similar conclusion, showing that species with high LMA values are more shade-tolerant than species with low LMA values. Like our model, they also model the carbon economy of a plant, balancing rates of carbon capture with carbon loss to calculate plant growth rate, and they also include an explicit assumption of a positive relationship between LMA and leaf lifespan.

Our analysis generalizes previous results that dealt only with soil fertility (Weng et al., 2017) or only with light environment (Sterck et al., 2006, Falster et al., 2018). Our analysis also expands previous studies and shows that a single mechanism can explain why low LMA should be favored in all kinds of high productivity environments while high LMA should be favored in all kinds of low productivity environments.

Another feature of our model is that it is simpler than previous similar models. Previous models contained assumptions about plant architecture (Sterck et al., 2006) or about allometric rules for biomass allocation between different tissues (Falster et al., 2018). The fact that we are able to recreate the results of more complicated models with our simple model shows that the mechanism at work here is very general, it does not rely on any specific assumptions about plant architecture or biomass allocation, and it works for all productivity gradients.

### An explanation of the mechanism that operates in our model

Plant carbon allocation to constructing new leaves can be seen as an investment that should be returned by new carbon sequestered by these new leaves. To better understand how different LMA values affect plant fitness, we examine the ratio of carbon fixed by a leaf over its entire lifespan to the construction cost of this leaf, which we define as the leaf ‘*lifetime return on investment’*. LMA is positively related to the total amount of carbon fixed by a leaf over its entire lifespan since higher LMA is correlated with longer leaf lifespan. LMA is also positively related to the construction cost of the leaf. Therefore, both the nominator and the denominator of the *lifetime return on investment* increase with LMA but at different rates.

All evidence show that leaf lifespan increases with LMA in a super-proportional manner (*α* > 1), that is, doubling of LMA increases leaf lifespan by more than twice. In fact, according to global trait databases, as LMA increases by a factor of 10, leaf lifespan increases by a factor of ~50 (Wright et al., 2004, Falster et al., 2012, Xu et al., 2017). As a result of this super-proportionality, the *lifetime return on investment* of high LMA leaves is higher than that of low LMA leaves (although see, Mediavilla and Escudero, 2003, Kikuzawa and Lechowicz, 2006, Ye et al., 2022), explaining the potential advantage of high LMA leaves.

Given the potential advantage of high LMA species as a result of their higher lifetime return on investment, we ask what is the advantage of plants with low LMA leaves in high productivity environments? we suggest that the explanation lies in the fact that plants with low LMA leaves have a higher rate of return on investment (as opposed to their lower lifetime return on investment) than plants with high LMA leaves. That is, it takes less time to return the cost of investment for low LMA leaves than for high LMA leaves. This allows reinvesting the assimilated carbon in the construction of new leaves, thus earning a compound interest on the investment (Westoby et al., 2000). Our model predicts that this compound interest is particularly valuable in high productivity “high interest” environments therefore providing an advantage for low LMA leaves under these conditions.

We analyzed the significance of the super-proportional relationship between LMA and lifespan by testing the hypothetical scenario of *α* ≤ 1, and showed that under such conditions the advantage of high LMA is lost and plants with low LMA leaves will always grow faster than plants with high LMA leaves. This is not surprising given that in the case of *α* = 1, the *lifetime return on investment* is equal for all plants regardless of their LMA, and if *α* < 1, the *lifetime return on investment* of plants with low LMA leaves is higher than that of those with high LMA. Furthermore, regardless of the value of *α*, plants with low LMA still retain a higher rate of return on investment compared to plants with high LMA.

Our model predicts that species with low LMA should have higher relative growth rates than species with high LMA under high productivity conditions. Indeed, LMA values were shown to be negatively correlated with relative growth rate under optimal growing conditions (Poorter and Remkes, 1990). Our model also predicts that plants with low LMA leaves will have a lower relative growth rate than plants with high LMA leaves under low productivity conditions. However, to the best of our knowledge there are no experiments that examined this hypothesis directly.

Another assumption of our model, in addition to the relationship between LMA and leaf lifespan, is that carbon acquisition is limited by leaf area. This is a natural assumption since photosynthesis is almost universally light limited (at least for parts of the day), except in extreme environmental conditions such as extremely dry conditions. In addition, increasing leaf area cannot increase carbon acquisition under closed canopy conditions, if there is no space to position new leaves such that they will receive enough light to be above the light compensation point. Our model will still apply if carbon acquisition is co-limited by leaf area and by additional factors (e.g., co-limitation by water and light availability).

## Conclusions

We presented a model that shows that the LMA value that optimizes growth rate decreases with increasing productivity, or more generally, with increasing net assimilation rate. Thus, our model provides a unified explanation for the observed negative correlations between plant LMA and habitat productivity. The central assumption that is required to explain why low LMA values would be favored in all kinds of high productivity environments is that leaf lifespan increases with LMA in a super-proportional manner. Our model sheds light on the mechanism relating LMA to plant fitness in different environments, thus contributing to greater mechanistic understanding of how plant traits influence plant fitness.

## Supplementary material – Deriving the analytic solution of the model

Here we show how we arrived at the analytic solution for the LMA value that optimizes relative growth rate. We derived equation 6 with respect to LMA to find the extremum points of the growth rate as a function of LMA. The derivative with respect to LMA is:

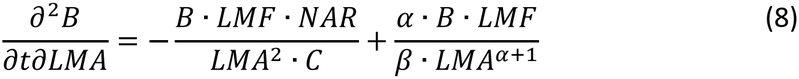

Equating the derivative to zero to find the LMA value that maximizes growth rate, we get:

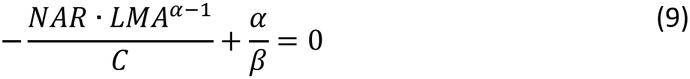

Therefore, the derivative equals zero when:

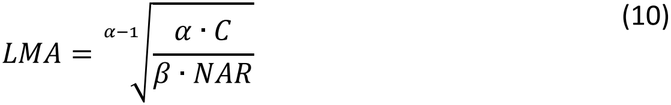

For the case of *α* > 1, which is most supported by empirical evidence, the extremum point above is a global maximum because the sign of the derivative changes from positive to negative at this point. Therefore, this extremum is also the LMA value that maximizes growth rate.

For the case of 0 < *α* < 1 the above extremum is a minimum rather than a maximum because the sign of the derivative changes from negative to positive at the extremum. Under this condition, the growth rate tends to infinity as LMA tends to zero.

For *α* = 1 we get that the derivative of the growth rate with respect to LMA is:

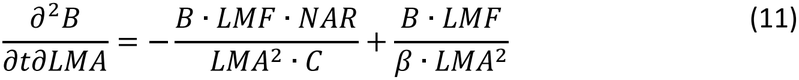

Therefore, when *α* = 1 there is no extremum (the derivative never equates to zero) and the growth rate is a monotonic (decreasing or increasing) function of LMA. The derivative is positive in the entire range when,

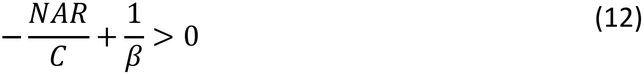

and then the growth rate is an increasing function of LMA and so the larger the LMA the higher the growth rate. However, in this case the growth rate of all the plants regardless of LMA is negative, and this case is not biologically relevant. Otherwise, the derivative is negative in the entire range when,

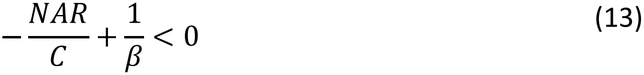

and then the growth rate is a decreasing function of LMA, so that the lower the LMA the higher the growth rate. In this case the relative growth rate is positive in some range of (low) LMA values. In the unique case of:

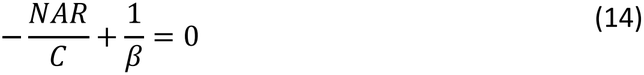

the derivative equals zero, and the growth rate does not depend on LMA.

To summarize, when *α* > 1 there is a single finite value of LMA that maximizes growth rate. The optimal value of LMA depends on the rate of net photosynthesis *NSR*, and on the values of other plant constants *c, α* and *β*. Otherwise, when 0 < *α* ≤ 1 no such finite value of LMA exists, and growth rate either keeps increasing as LMA tends to infinity or keeps increasing as LMA tends to zero or is constant.

### Alternative version of the model where leaf mass fraction (LMF) depends on LMA

In order to check if the assumption that LMF is independent of LMA is important for the qualitative behavior of the model, we examined an alternative version of the model. In this alternative version, we assume that LMF is dependent on LMA and that dependency is given by:

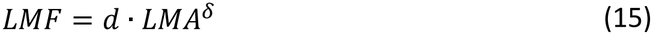

Then, substituting this dependency in equation (6) we get:

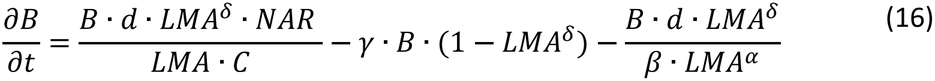

We examined the behavior of this equation using a numerical solution along the range of naturally existing LMA values and with different parameter sets for the values of *d* and *δ*. The results are shown in figures 3-5.

**Figure 3:**
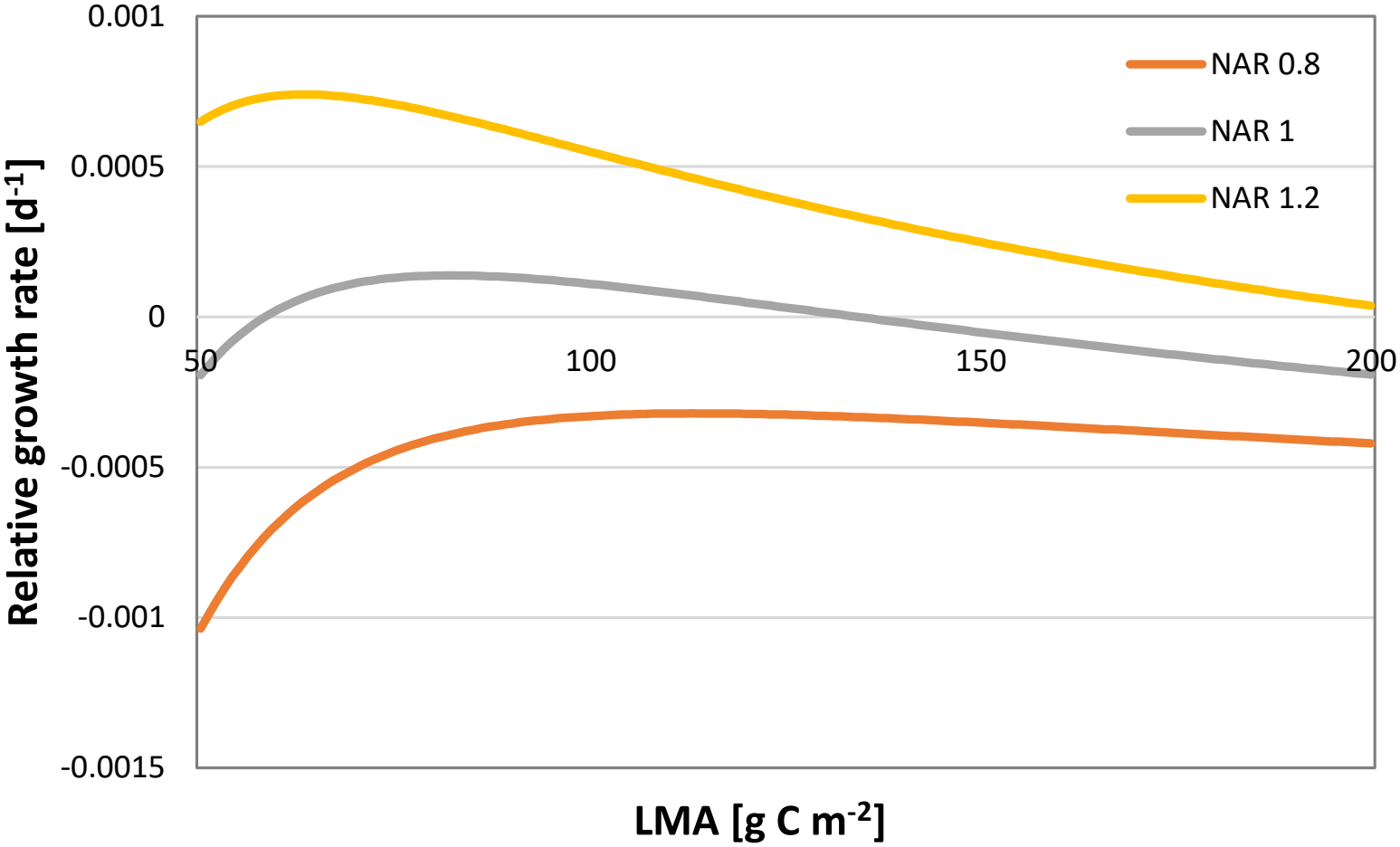
Daily rate of relative growth as a function of leaf mass per area (LMA) with three different values of the net assimilation rate (NAR). The values of the parameters are as follows: *α* = 1.71, *β* = 0.092923, c = 1.2 and *γ* = 0.001369. The values of *α* and *β* are based on Wright et al., 2004, the value of c is a typical value for this trait and *γ* was calculated as the inverse of a turnover time of two years (730 days) which is a reasonable turnover time for non-leaf tissue. The values for the scaling of LMF with LMA are d = 0.2 and δ = 0.06.

**Figure 4:**
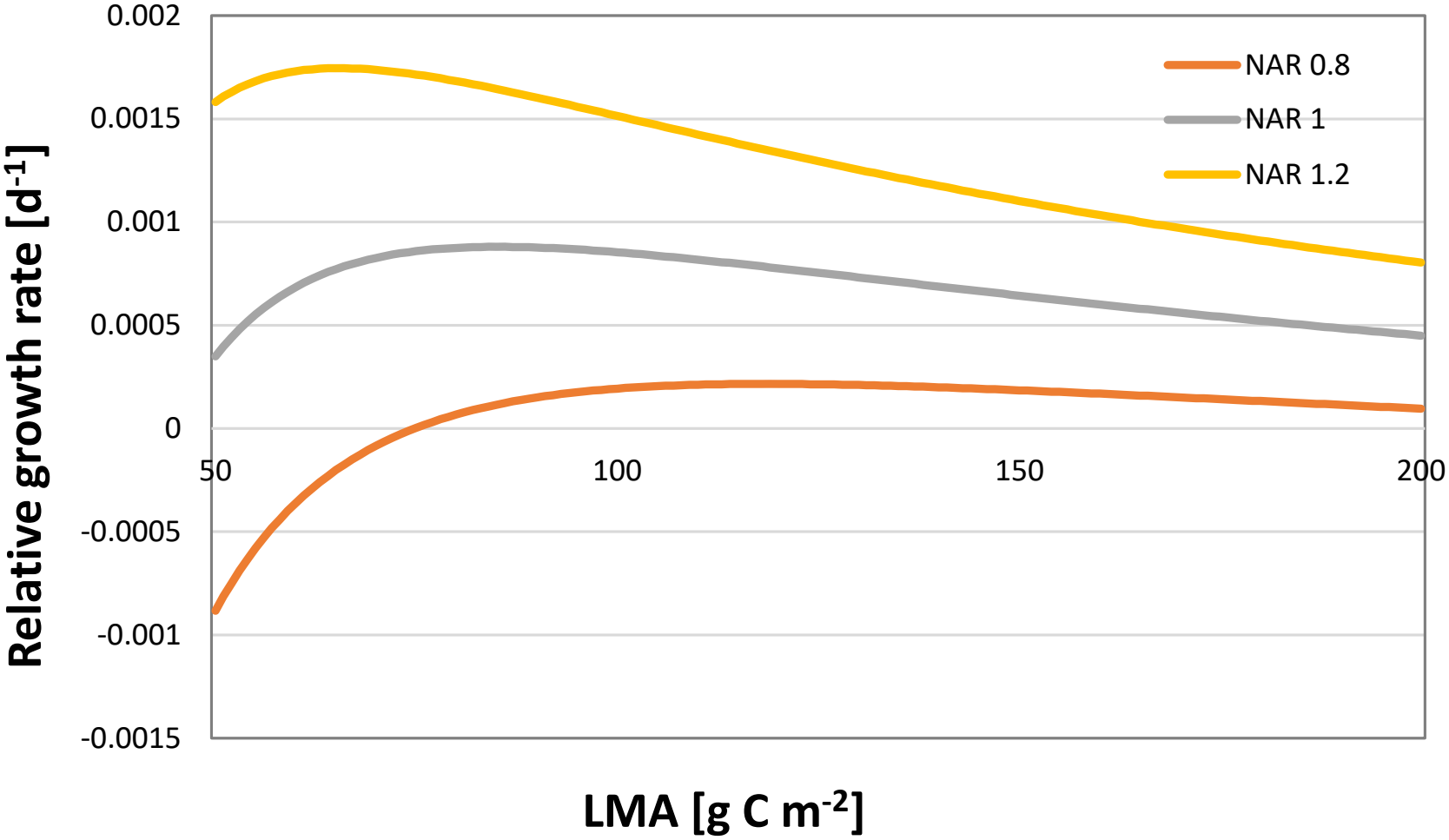
Same as figure 3 except that the values for the scaling of LMF with LMA are d = 0.25 and δ = 1.

**Figure 5:**
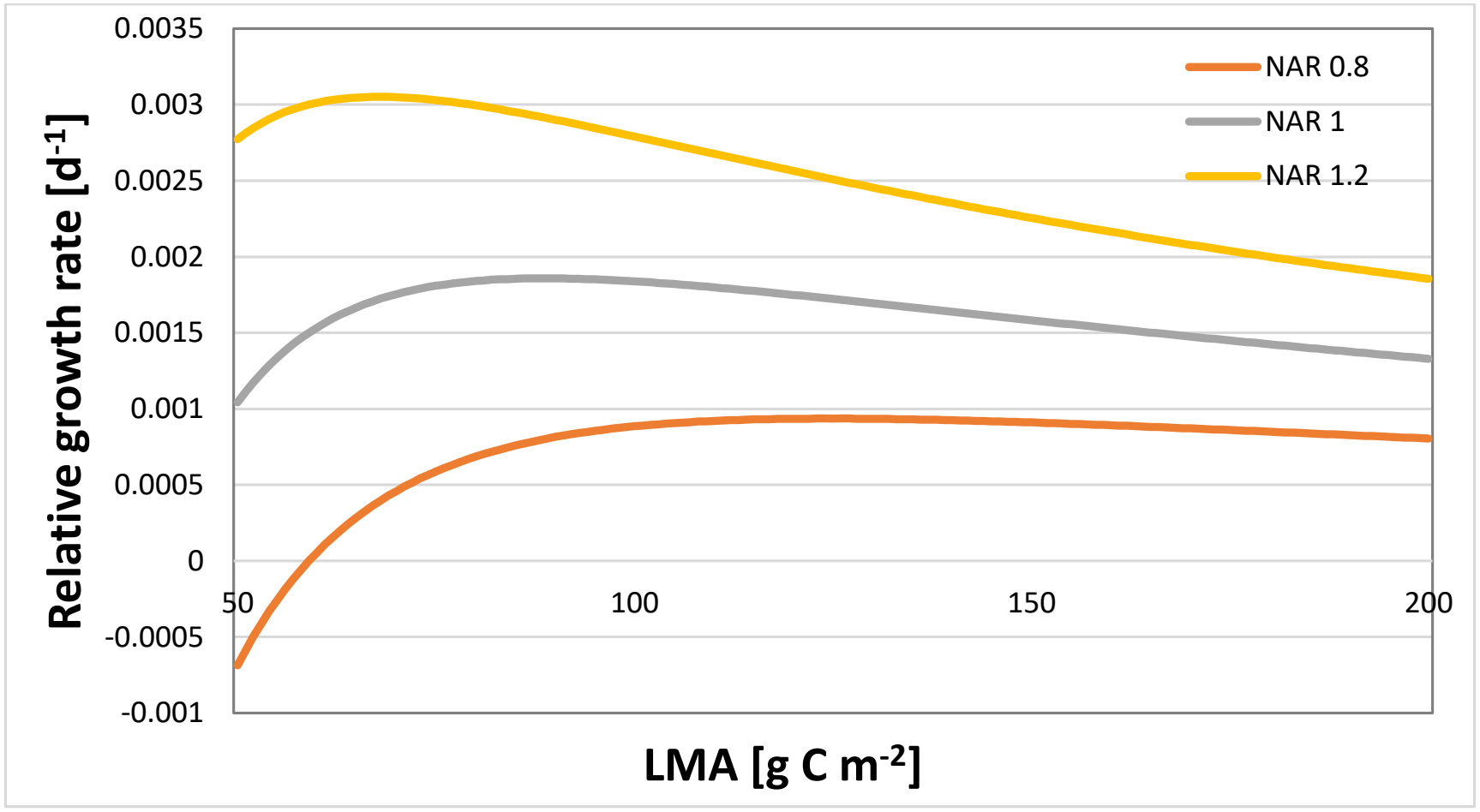
Same as figure 3 except that the values for the scaling of LMF with LMA are d = 0.3 and δ = 0.14.

The analyses show that the qualitative behavior of the model remains unchanged relative to the original model. There is a single value of LMA that optimizes growth rate, and the optimal LMA value decreases with increasing NAR. For all the parameter sets we have tested the values of LMF are within a normal range for this trait and the relative growth rate that is achieved with the optimal value of LMA is positive with only one exception that occurs with the lowest values of *d, δ* and NAR (figure 3).

